# Microarray screening of differentially expressed genes after up-regulating miR-205 or down-regulating miR-141 in cervical cancer cells

**DOI:** 10.1101/618041

**Authors:** Xingyu Fang, Tingting Yao

**Affiliations:** Department of Gynecological Oncology, Sun Yat‐sen Memorial Hospital, Sun Yat‐sen University, Guangzhou, China; Guangdong Provincial Key Laboratory of Malignant Tumor Epigenetics and Gene Regulation, Sun Yat-Sen Memorial Hospital, Sun Yat-Sen University, Guangzhou, China

**Keywords:** Gene expression profiling, cervical cancer, miRNA

## Abstract

Cervical cancer is one of the most common gynecological malignancies. However,studies on the expression and molecular mechanism of miR-205 and miR-141 in CC are insufficient recently. Expression profile microarray with 21329 Oligo DNA were used to detect the expression of mRNAs in miR-205 up-regulated or miR-141 down-regulated HeLa and SiHa cells and mRNAs in normal HeLa and SiHa cells. Gene Ontology (GO), Kyoto Encyclopedia of Genes and Genomes (KEGG) pathway were performed to assess the potential pathways of miR-205 in SiHa cell.Compared with normal HeLa cell, there were 38 differentially expressed genes (DEGs) in miR-205 up-regulated HeLa cell. Nine were up-regulation genes and 29 were down-regulation genes. There were 23 DEGs in miR-141 down-regulated HeLa cell. One was up-regulated and 22 were down-regulated. Compared with normal SiHa cell, there were 128 DEGs in miR-205 up-regulated SiHa cell. One hundred and three were up-regulation genes and 25 were down-regulation genes. There were 80 DEGs in miR-141 down-regulated SiHa cell. Forty two were up-regulation genes and 28 were down-regulation genes. For miR-205 up-regulated SiHa cell, GO outcome showed that “ubiquitin-protein ligase activity”, “MAP kinase phosphatase activity”, were the most enriched terms (*P* < 0.05). And in KEGG analysis, “Cell cycle” was notably enriched, and Smad4 in this pathway was up-regulated (*P* < 0.05). Expression profile microarray technology can effectively screen out DEGs in cervical cancer cells after up-regulating miR-205 or down-regulating miR-141. Which may enable us to understand the pathogenesis and lay an important foundation for the prevention and treatment of cervical cancer.

## Introduction

Cervical cancer is one of the most common gynecological malignancies and the fourth leading cause of cancer-related death worldwide. There were 527,600 new cases of cervical cancer and 265,700 deaths worldwide in 2012.(1) It is the most common cancer of women in many low-income countries.(2) Persistent infection of human papillomavirus (HPV) type 16 and 18 are the major risk factors for cervical cancer. In HPV high-risk cervical cancer cases, HPV16 and 18 phenotypes account for 70%.(3) After HPV DNA integrated into the host DNA, *E6* and *E7* genes are expressed. *E6* induces the degradation of p53, stimulates telomerase activity, and induces centrosome distortion and instability of the chromosome.(4) The binding of *E7* protein with human pRb and E2F transcription factor mediates cell transformation, leading to the separation of pRb and E2F transcription factor and making cells enter S phase of cell cycle early.(5) The occurrence of cervical cancer is a multi-factor, multi-stage and multi-gene process, involving a large number of structural changes and abnormal expression of tumor-related genes. It is particularly important to study the molecular mechanism of the oncogenesis and development of cervical cancer and to select effective therapeutic targets.

MicroRNAs (miRNAs), usually have 18-25 nucleotides, are a kind of small highly conserved non-coding RNA and are endogenous expression in many species. Most mature miRNAs inhibit gene expression by cutting target messenger RNA(mRNAs) or inhibiting mRNAs’ translation by binding to the 3 ’-UTRs of mRNAs.(6) More and more reports have shown that miRNA plays an important role in the proliferation, apoptosis, invasion and migration of tumor cells. Abnormal expression of miRNA can be used for tumor classification, diagnosis and prognosis.(7) The carcinogenic or cancerocidal effects of miRNAs in the tumorigenesis and development of cervical cancer depend on the biological effects of their target genes.(8)

Microarray-based gene expression profiling technology has been applied to analyze the regulatory changes and expression differences of genes in tissues and cells, considerably facilitating the exploration of molecular mechanisms of tumor development. However, reports on the expression and molecular mechanism of miR-205 and miR-141 in CC are insufficient recently. In this study, microarray-based gene expression profiling technology was used to detect DEGs after up-regulation of miR-205 or down-regulation of miR-141 in SiHa and HeLa cells. Then screening out the gene groups related to the occurrence and development of cervical cancer, studying their functions, exploring the relationship between DEGs and the tumorigenesis and progression of cervical cancer in order to provide evidence for the selection of therapeutic targets for cervical cancer.

## Materials and methods

### Cell culture

The *human* cervical cancer cell lines SiHa and HeLa were purchased from American Type Culture Collection (ATCC, Manassas, VA) and cultured at 37°C in a humidified 5% CO2 atmosphere in DMEM medium (Gibco; Thermo Fisher Scientific, Inc.) with 10% fetal calf serum (Gibco; Thermo Fisher Scientific, Inc.),100 IU/ml penicillin G, and 100 mg/ml streptomycin sulfate (Sigma‑Aldrich; Merck KGaA, Darmstadt, Germany).

### SiRNA transfection

Synthetic small interfering RNA (siRNA) used to knockdown miR-141 or overexpress miR-205 were purchased from GenePharma (Suzhou, China). siRNAs were transfected into cells cultured in 10cm culture dish using Lipofectamine 3000 (Invitrogen) according to the manufacturer’s instructions. Total RNA was extracted using the Trizol reagent (Invitrogen, Carsibad, CA, USA) according to the manufacturer’s protocol 48h after transfection.

### The main experimental instruments

The 22K *Human* Genome Array is provided by the CaptialBio Microarray service (CapitalBio, Beijing, China), according to the manufacturer’s instructions, and 21329 Oligo DNA of this genome array were from Human Genome Oligo Set Version 2.1, Operon company. The other 193 Oligo DNA were synthesized by CapitalBio company. The scanner is LuxScan 10KA dual-channel laser scanner (CapitalBio, Beijing, China).The analysis image software is LuxScan 3.0 (CapitalBio, Beijing, China).

### Preparation method of chip

For each RNA array experiment, we absorbed 1ul RNA to test the concentration and purity by spectrophotometer at 260nm/180nm, then electrophoresed RNA on gel containing formaldehyde for quality control [Fig1].

**Fig1:**
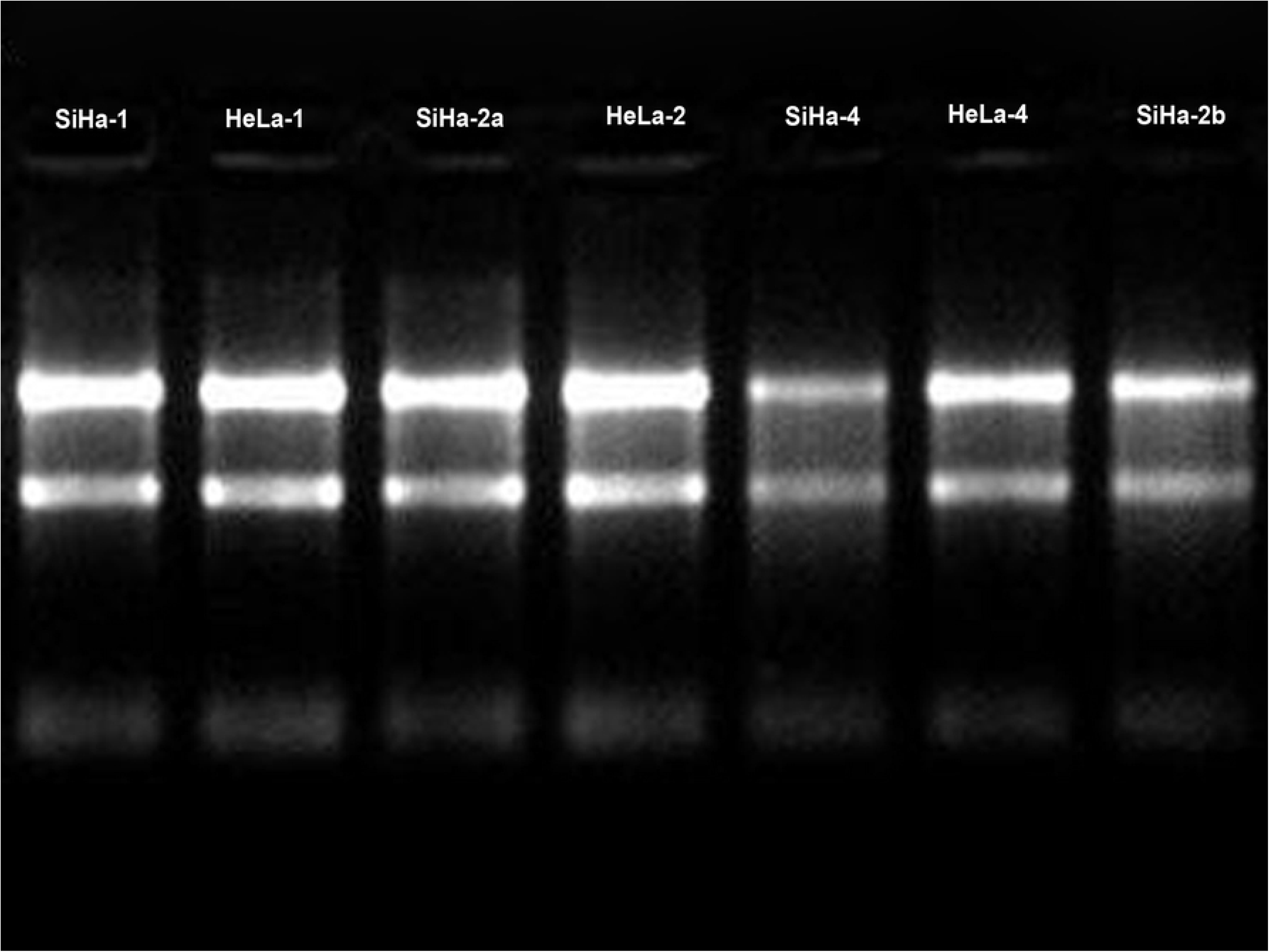
RNA formaldehyde modified gel electrophoresis. From left to right are SiHa-1, HeLa-1, SiHa-2a, HeLa-2, SiHa-4, HeLa-4, SiHa-2b. The bands are clear, and the brightness of 28S is greater than that of 18S.

For each array experiment, we extracted 4ul of total RNA, added to 50ul reverse transcription reaction system. The samples were fluorescently labeled with cRNA, purified and dried, and then used for chip hybridization. mRNA of HeLa-1,HeLa-2,SiHa-1,SiHa-2 was labeled with cy5-2 dUTP. mRNA of HeLa-4,SiHa-4 was labeled with cy3-2 dUTP. Then we dissolved samples respectively in 20ul hybridization buffer containing 5× sodium chloride-sodium citrate Buffer (SSC), 0.2% sodium dodecyl sulfate (SDS). Hybridization was performed at 42°C overnight. Gene chip and hybridization probes were respectively degenerated 5 min in 95 °c water bath, then added a probe immediately on the chip. We used cover glass sealing piece, placed pieces inside the 60 °c capsule and hybridized 16 h. The mRNA array was washed with 2×SSC and 0.2%SDS solution, 0.1%SSC and 0.2%SDS solution, 0.1%SSC solution respectively at 42°C for 10 min, then dried at room temperature. The array was scanned by a Dual channel laser scanner (LuxScanTM 10K/A, CaptialBio). LuxScan3.0 image analysis software (CapitalBio) was used to analyze the chip image and convert the image signal into digital signal. The Lowess method is used to normalize the data on the chip.

### Gene expression analysis, gene ontology, pathway analysis

mRNAs with a IV signal value ≥1500, and at least a 2-fold increase or a 0.5-fold decrease in mRNA expression were considered to have significant changes, including up-regulation and down-regulation. We selected DEGs between SiHa-1 and SiHa-4 to conduct functional enrichment analysis using Gene Ontology (GO) based on the MAS (molecule annotation system), and perform pathway analysis using Kyoto Encyclopedia of Genes and Genomes (KEGG) database. (http://www.kegg.jp/)

### Statistical analysis

We use Wilcoxon signed-rank test to determine the statistical difference between two groups. And the hypergeometric distribution test was used to perform GO and pathway enrichment analysis. *P* value of less than 0.05 was considered statistically significant in these texts. Statistical analysis was executed using SPSS software version 20.

## Results

The total RNA extracted from all samples ranged from 16.35 to 18.34 ug, and the absorbance of 260nm/280nm ranged from 2.09 to 2.10. The concentration and purity of all RNA samples were good. RNA electrophoresis band [Fig1] showed the quality is up to standard and the quantity is enough.

In all chip images, the signal of positive controls (HEX), external markers (eight intergenic sequences of yeast) were normal, and the signal of negative controls (12 artificially synthesized 70 mer Oligo DNA with no homology with human genes and 50%DMSO) were negative. The repeatability of house-keeping gene was good and the Ratio CV was no more than 0.3, the leakage rate did not exceed 3‰.

According to the statistical selection criteria, our results showed that there were 38 DEGs between HeLa-1 and HeLa-4 [Table1]. Nine were up-regulation genes and 29 were down-regulation genes. There were 23 DEGs between HeLa-2 and HeLa-4 [Table2]. One was up-regulated and 22 were down-regulated. There were 128 DEGs between SiHa-1 and SiHa-4 [Table3]. Of these, 103 were up-regulation genes and 25 were down-regulation genes. There were 80 DEGs between SiHa-2 and SiHa-4 [Table4]. Fifty-two were up-regulation genes and 28 were down-regulation genes. The outcome of the hybridization scatter plot were the same [Fig2].

**Fig2:**
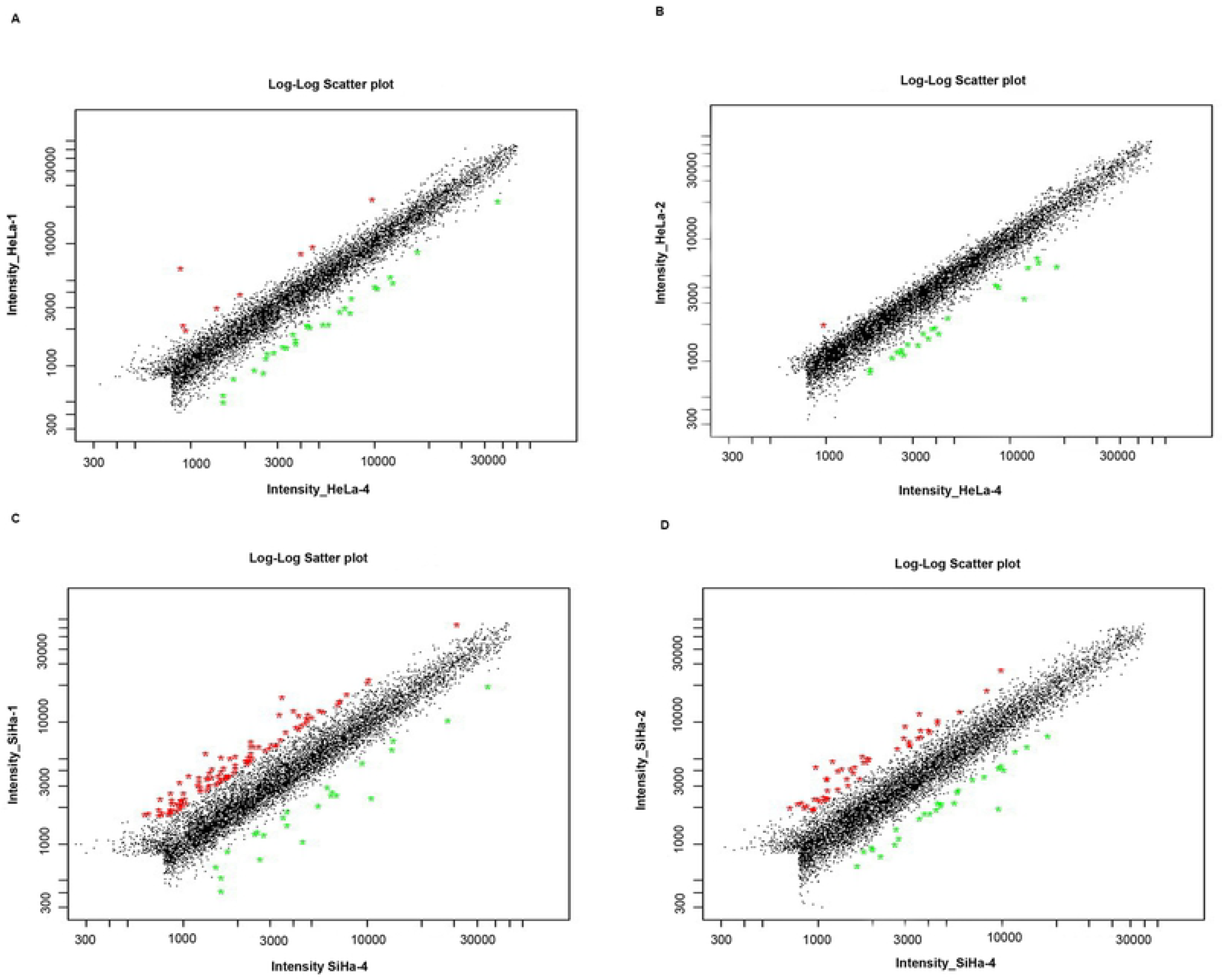
(A) Hybridization scatter plot of HeLa-1 and HeLa-4.(B) Hybridization scatter plot of HeLa-2 and HeLa-4.(C) Hybridization scatter plot of SiHa-1 and SiHa-4.(D) Hybridization scatter plot of SiHa-2 and SiHa-4. The X axis and Y axis coordinates respectively represented the fluorescence signal strength value of two samples, each point in the graph represent one gene. Red and green points respectively represented the ratio of Y/X ≥2 or ≤0.5, black points represented the ratio of Y/X were between 0.5 and 2.0.

**Table 1:** Part of the differentially expressed genes in miR-205 up-regulated HeLa cell.

| Gene name of the<br>Chip <sup>a</sup> | Gene name <sup>b</sup> | Up-regulation rate <sup>c</sup> | Down-regulation rate<br><sup>c</sup> |
| --- | --- | --- | --- |
| DOCK3 | - | 18.8572 |  |
| KIAA1233 | ADAMTSL3 | 7.1471 |  |
| ODC1 | ODC1 | 2.3646 |  |
| HT007 | - | 2.1446 |  |
| COL4A3BP | COL4A3BP | 2.0514 |  |
| PPID | PPID | 2.0482 |  |
| FLJ21802 | - | 2.0426 |  |
| GRWD | GRWD1 | 2.0381 |  |
| CASP1 | CASP1 |  | 0.3754 |
| IGFBP1 | IGFBP1 |  | 0.3854 |
| CHRD | CHRD |  | 0.4046 |
| FLJ23058 | Q9P281 |  | 0.407 |
| MMD | MMD2 |  | 0.4102 |
| PROS1 | PROS1 |  | 0.4129 |
| WFDC1 | WFDC1 |  | 0.4148 |
| DUSP5 | DUSP5 |  | 0.4265 |
| SERPINE1 | SERPINE1 |  | 0.4311 |
| BHLHB2 | BHLHB2 |  | 0.4341 |
| BF | BF |  | 0.4363 |
| ATIP1 | MTUS1 |  | 0.4463 |
| IL11RA | IL11RA |  | 0.4464 |
| KIAA0603 | TBC1D4 |  | 0.4544 |

a The gene name defined by chip
b Official gene name
c The ratio of Cy5/Cy3. Cy5 is the signal value of the gene detected in HeLa-1. Cy3 is the signal value of the gene detected in HeLa-4

**Table2:** Part of the differentially expressed genes in miR-141 down-regulated HeLa cell.

| Gene name of the Chip<br>a | Gene name <sup>b</sup> | Up-regulation rate <sup>c</sup> | Down-regulation<br>rate <sup>c</sup> |
| --- | --- | --- | --- |
| IGFBP3 | IGFBP3 |  | 0.2691 |
| CREM | - |  | 0.3295 |
| COL15A1 | - |  | 0.398 |
| BF | BF |  | 0.4159 |
| GCLC | - |  | 0.4249 |
| UBD | UBD |  | 0.4394 |
| IL6 | IL6 |  | 0.4556 |
| JDP1 | DNAJC12 |  | 0.4556 |
| ZNF148 | - |  | 0.4593 |
| ETV4 | ETV4 |  | 0.4601 |
| LOC58489 | Q9NPM1 |  | 0.4637 |
| CASP8AP2 | CASP8AP2 |  | 0.4716 |
| HBP17 | FGFBP1 |  | 0.4746 |
| ATP6C | ATP6V1C1 |  | 0.4748 |

a The gene name defined by chip
b Official gene name
c The ratio of Cy5/Cy3.Cy5 is the signal value of the gene detected in HeLa-2.Cy3 is the signal value of the gene detected in HeLa-4

**Table3:**
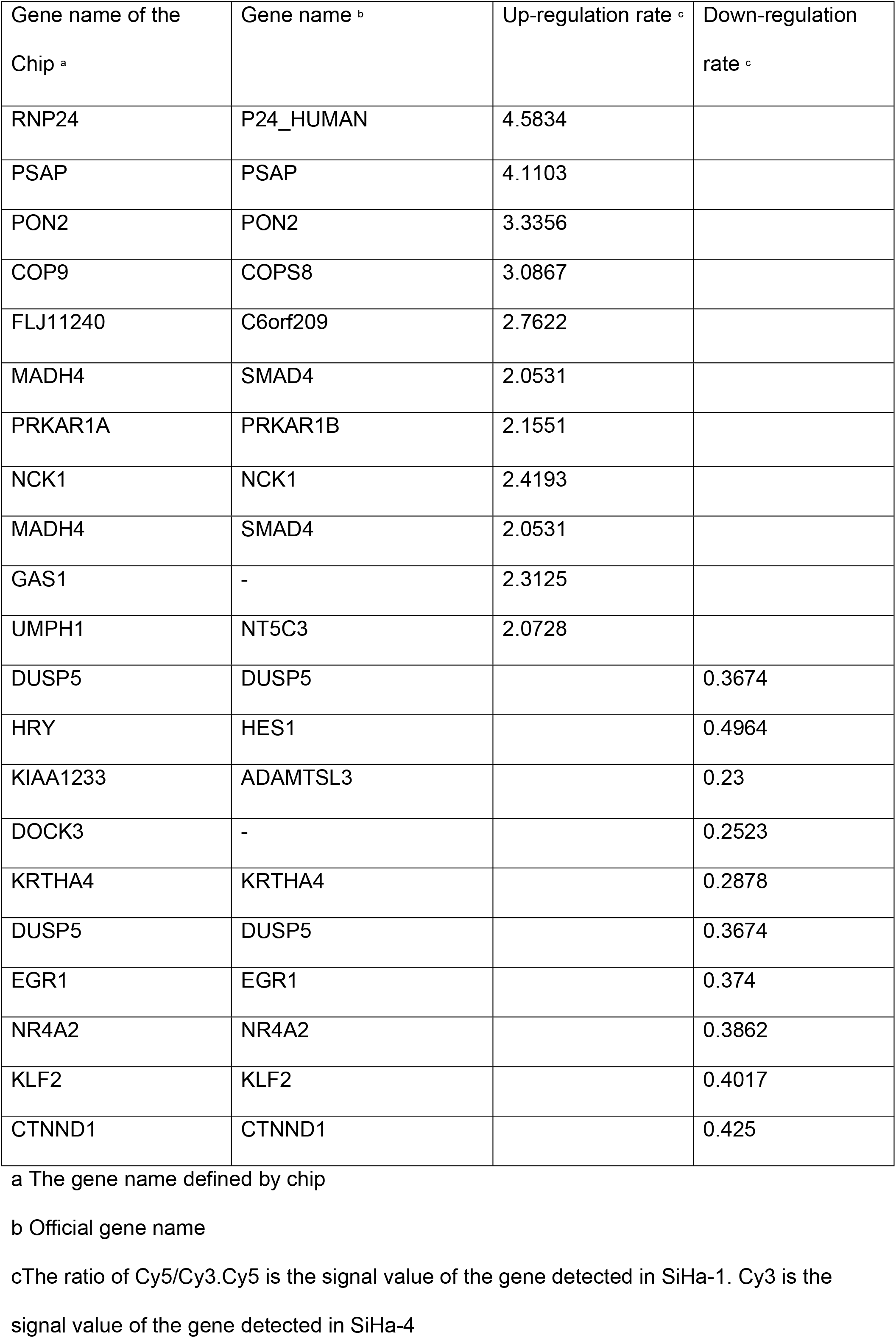
Part of the differentially expressed genes in miR-205 up-regulated SiHa cell.

**Table4:**
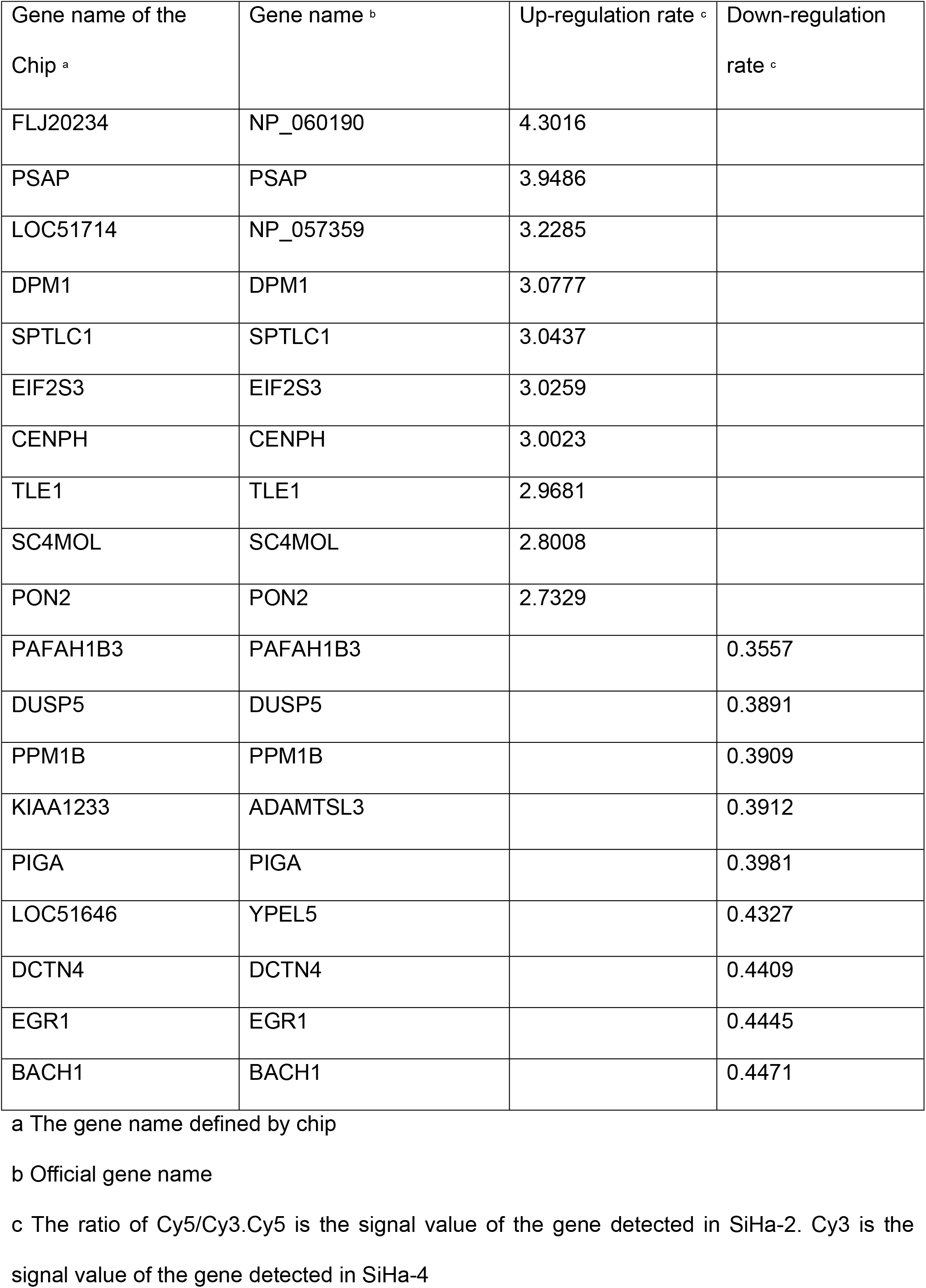
Part of the differentially expressed genes in miR-141 down-regulated SiHa cell.

| Gene name of the<br>Chip <sup>a</sup> | Gene name <sup>b</sup> | Up-regulation rate <sup>c</sup> | Down-regulation<br>rate <sup>c</sup> |
| --- | --- | --- | --- |
| FLJ20234 | NP_060190 | 4.3016 |  |
| PSAP | PSAP | 3.9486 |  |
| LOC51714 | NP_057359 | 3.2285 |  |
| DPM1 | DPM1 | 3.0777 |  |
| SPTLC1 | SPTLC1 | 3.0437 |  |
| EIF2S3 | EIF2S3 | 3.0259 |  |
| CENPH | CENPH | 3.0023 |  |
| TLE1 | TLE1 | 2.9681 |  |
| SC4MOL | SC4MOL | 2.8008 |  |
| PON2 | PON2 | 2.7329 |  |
| PAFAH1B3 | PAFAH1B3 |  | 0.3557 |
| DUSP5 | DUSP5 |  | 0.3891 |
| PPM1B | PPM1B |  | 0.3909 |
| KIAA1233 | ADAMTSL3 |  | 0.3912 |
| PIGA | PIGA |  | 0.3981 |
| LOC51646 | YPEL5 |  | 0.4327 |
| DCTN4 | DCTN4 |  | 0.4409 |
| EGR1 | EGR1 |  | 0.4445 |
| BACH1 | BACH1 |  | 0.4471 |

a The gene name defined by chip
b Official gene name
c The ratio of Cy5/Cy3. Cy5 is the signal value of the gene detected in SiHa-2. Cy3 is the signal value of the gene detected in SiHa-4

The function classification of DEGs that was identified between SiHa-1 and SiHa-4 was conducted with GO enrichment analysis. These genes were cataloged according to biological processes (BPs), molecular functions (MFs), and cellular components (CCs) based on GO database, and we showed the most likely (with the lowest *P* value) relevant 10 Go terms [Fig3].We found that for molecular function, the DEGs mainly involved in the ubiquitin-protein ligase activity, MAP kinase phosphatase activity, transcription factor activity. For biological processes, the DEGs were mostly enriched in ubiquitin cycle, regulation of transcription, DNA-dependent, cell cycle arrest. For cellular Component, the DEGs were distributed mainly in nucleus and mitochondrion.

**Fig3:**
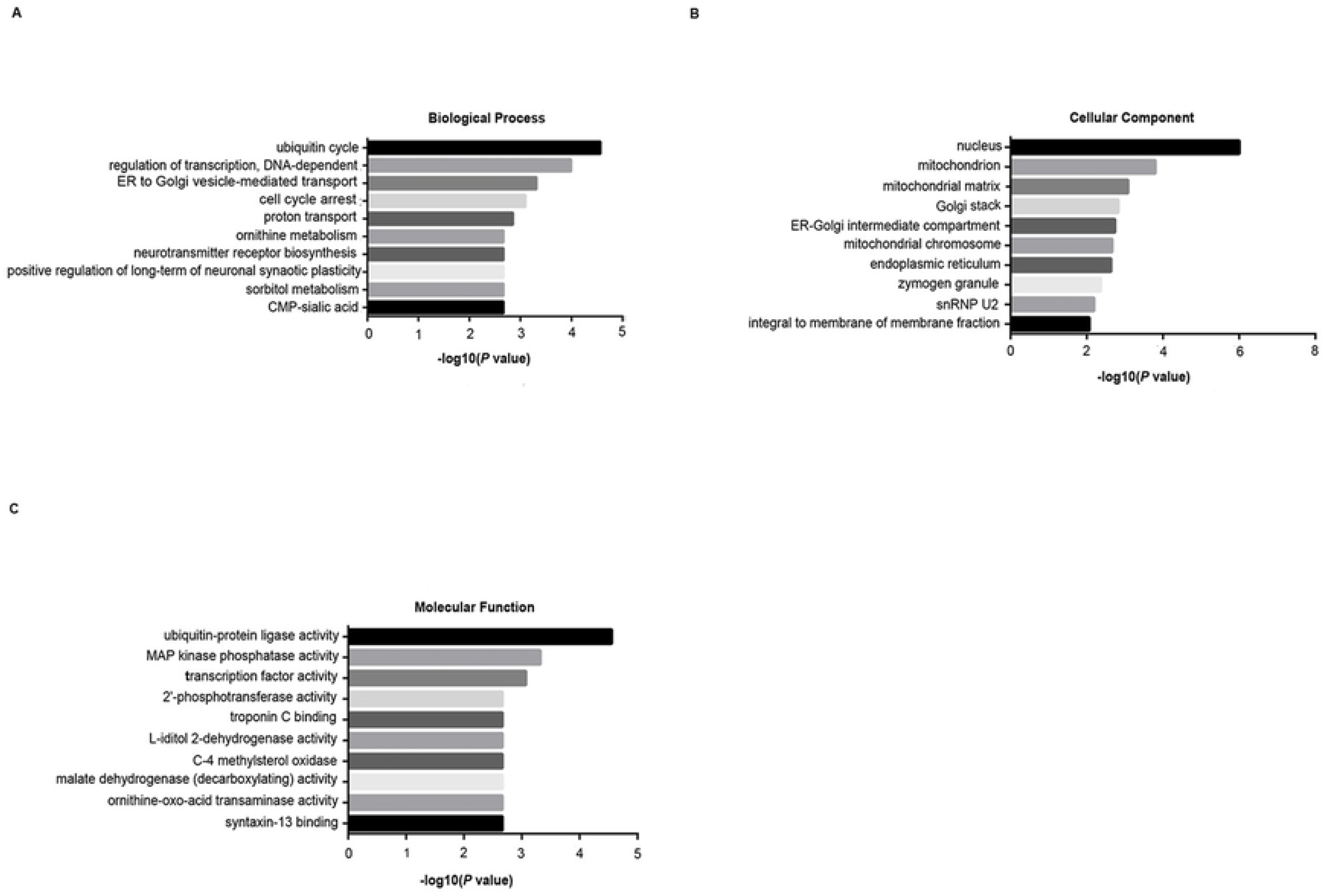
GO enrichment analysis of miR-205 up-regulated SiHa cell, in terms of (A) biological process, (B) cellular component and (C) molecular function.

The concentration trend of the DEGs between SiHa-1 and SiHa-4 were further defined based on KEGG pathway analysis. Eight pathways with *P*<0.05 were listed in Table5. The concentration trend was inversely proportional to the *P* value. The most likely involved pathways were cell cycle, sphingolipid metabolism, ubiquitin-mediated proteolysis, and pyrimidine metabolism. And the DEGs in these signaling pathways were shown in Fig 4.

**Fig4:**
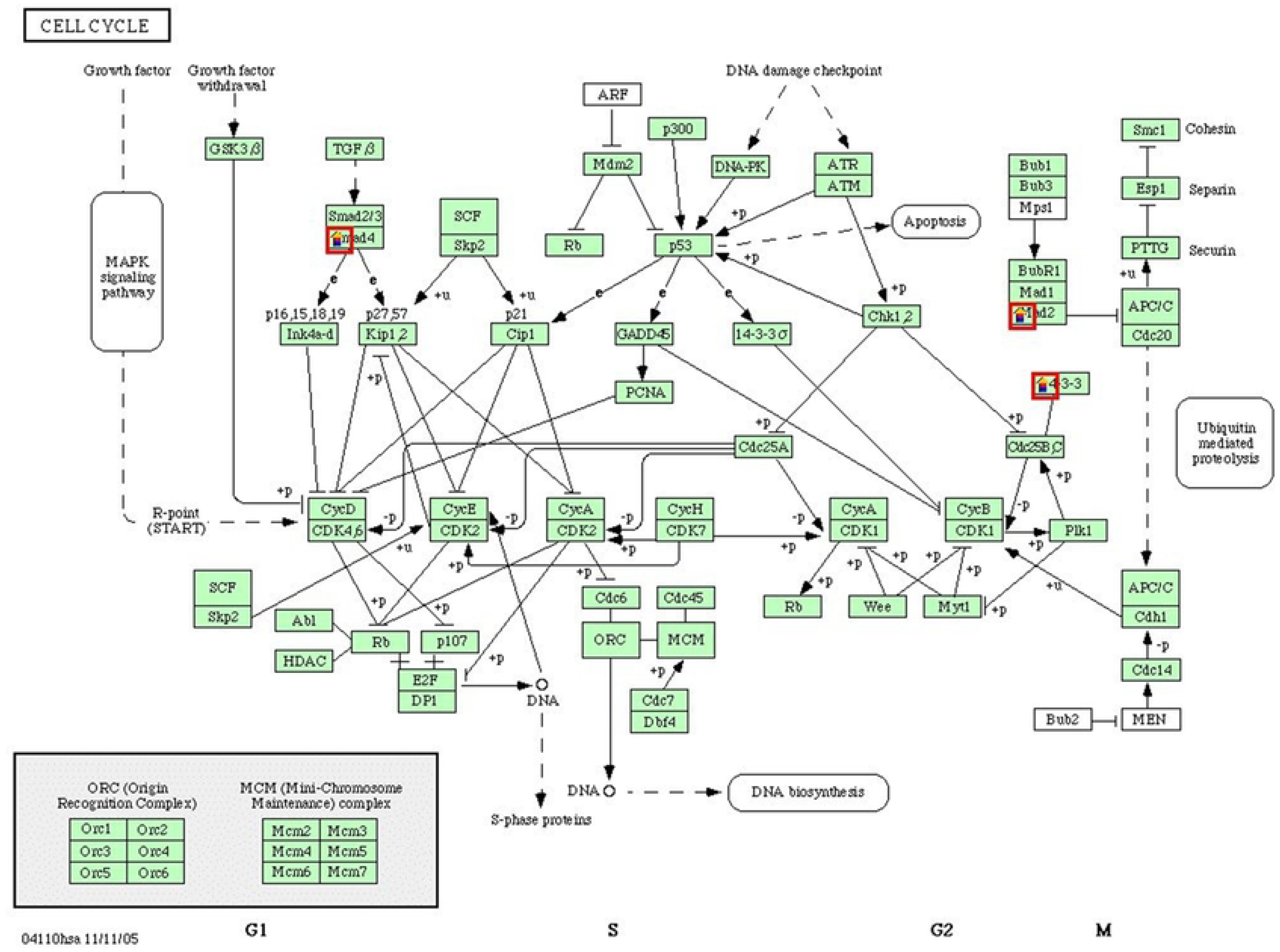
The differentially expressed genes and KEGG pathways of miR-205 up-regulated SiHa cell were predicted by KEGG.

**Table5:** The pathways of differentially expressed genes in miR-205 up-regulated SiHa cell.

| Pathway Name <sup>a</sup> | Gene name <sup>b</sup> | *P* value |
| --- | --- | --- |
| Cell cycle | SMAD4,MAD2L1,YWHAG | 0.004407 |
| Sphingolipid metabolism | DEGS1,SPTLC1 | 0.007082 |
| Nicotinate and nicotinamide metabolism | NT5C1B,NT5C3 | 0.007082 |
| Ubiquitin mediated proteolysis | UBE2D3,UBE2D2 | 0.007735 |
| Valine, leucine and isoleucine degradation | NAT5,ACADM | 0.010991 |
| Pyrimidine metabolism | NT5C1B、 NT5C3 | 0.030525 |
| Bisphenol A degradation | PON2 | 0.042995 |
| Ethylbenzene degradation | NAT5 | 0.048587 |

a The pathway defined based on KEGG pathway analysis
b The gene enriched in the signal pathways

## Discussion

Tumorigenesis is a complex process in which multiple genes are involved. In addition to the expression changes of gene related to cell proliferation, apoptosis and transcription, the changes connected with some biological processes in cells also play a part.

Recent years, miRNAs have been considered as key regulatory molecules of multiple signaling pathways including tumor development, differentiation, proliferation, apoptosis and invasion.(9–10) The increasing maturity of gene expression profile microarray technology provides the foundation for finding out the mechanism of tumorigenesis on the whole. In our research, we up-regulated miR-205 in SiHa cell, discovered that the DEGs covered multiple biological processes, mainly including ubiquitin cycle, DNA-dependent transcriptional regulation, cell cycle arrest and transcription. These genes also involved in multiple signaling pathways such as cell cycle and tumor formation. Therefore, we believe that *miR-205*, a target for the regulation of the occurrence and development of cervical cancer, can provide better support and help for the treatment of cervical cancer by regulating its expression.

MiR-205 is located in chromosome 1q32.2. Low expression of miR-205 have been found in liver cancer, prostate cancer, breast cancer, renal cell cancer and so on.(11–13) While in ovarian cancer, miR-205 is highly expressed.(14) Therefore, we supposed that miR-205 exerts different effects on diverse cancers. Pang, h. and x. Yue’s et al. found that the ectopic expression of miR-205 inhibited the proliferation of CC cells and promoted apoptosis. Suggesting that miR-205 may be an anti-oncogene for cervical cancer.(15)

Our study found that after up-regulating miR-205, the expression of Smad4 in SiHa cells increased. Smad proteins are found in the cytoplasm and are signaling transduction elements of the transforming growth factor-β (TGF-β) superfamily.(16) Transforming growth factor-β (TGF-β) family members exert their function via specific type I and type II serine/threonine kinase receptors and intracellular Smad transcription factors.(17) The SMAD family can be divided into three categories: receptor-regulated Smad (R-Smad); Co-mediator Smad (Co-Smad); and the inhibitory Smad (I-Smad).Smad4 is the only Co-Smad. R-smads (Smad1, 2, 3, 5, 8) are directly phosphorylated and activated by type I receptor kinase and undergo homotrimerization and formation of heteromeric complexes with *Smad4*. Then, activated Smad compound are transferred to the nucleus, and regulate the transcription of target gene together with other nuclear coenzyme factors.(18) The absence of Smad4 plays an important role in skin and upper gastrointestinal squamous cell carcinoma and gastrointestinal adenocarcinoma.(17) Baldus, S.E et al. analyzed the expression of Smad4 in 13 cervical cancer cell lines, 40 precancerous lesions and 41 primary squamous cell carcinoma specimens, found that the loss of Smad4 may play an significant part in the occurrence of cervical cancer.(19)

More and more evidence indicates that microRNAs (MiRNAs) exert an enormous function on TGF-β/smad pathway-induced tumor suppression.(20–22) Members of the TGF-β family regulate a variety of biological processes, including morphogenesis, embryonic development, adult stem cell differentiation, immune regulation, wound healing, inflammation, and cancer. In the early stage of tumorigenesis, TGF-β can inhibit the proliferation but promote apoptosis of epithelial cells, endothelial cells and hematopoietic cells. In advanced tumor, TGF-β can promote the migration and invasion of cancer cells, facilitate angiogenesis and induce immunosuppression, the increase of TGF-β is related to the increased aggression of tumor and poor prognosis of patients.(23)

Smad4 protein significantly affects tumor angiogenesis, tumor proliferation, invasion, metastasis and other biological behavior mainly through TGF-β signal pathway. In pancreatic cancer, TGF-β inhibits cell growth and proliferation by blocking mitogenic growth signals through Smad4. It also induces programmed cell death (PCD) or apoptosis of pancreatic cells by regulating the TIEG, a zinc-finger gene transducted by TGF-β/Smad4 signaling, and takes part in tumor inhibition.(24) Smad proteins regulate the TGF-β signaling pathway, and the absence of the tumor suppressor. Smad4 is one of the molecular mechanisms leading to TGF-β resistance. Zhu, H et al. summarized the expression levels of TGF-β in cervical intraepithelial neoplasia (CIN), cervical cancer and normal tissues, concluded that TGF-β1 functions as a tumor inhibitor in precancerous lesions of cervical cancer and early cervical cancer, while as a tumor promotor in advanced tumors.(25)

The expression proteins of HPV16 push forward an immense influence on the regulation of TGF-β pathway through Smads protein. Among them, HPV-16 E7 protein is mainly related to immune stress and the promotion of tumor cell proliferation.(26) Lee, D.K. et al. demonstrated that E7 protein can bind to Smad2, Smad3 and Smad4, and block the interaction of Smad complex with the Smad DNA binding element, CAGA. Suggesting that E7 oncoprotein inhibits the signaling transduction of TGF-β by inhibiting Smad DNA binding activity.(27) The expression of HPV 16 E5 can attenuate the TGF-β/Smad signaling, and the loss of signal transduction can lead to the imbalance of epithelial homeostasis in the early stage of viral infection, which may be an crucial mechanism to promote the occurrence of HPV mediated cervical cancer.(28) Peralta-zaragoza et al. found that E6 and E7 oncoproteins can regulate the expression of TGF-β1 through increasing the activity of TGF-β1 promoter. Which reveals the contribution of HPV-16 to TGF-β1 gene expression in cervical cancer.(29)

TGF-β-induced growth responses differ between SiHa and HeLa cells. In SiHa cells, TGF-β-mediated growth inhibition depends on the nuclear accumulation of activated Smads. In HeLa cells, the cell cycle progression induced by TGF-β is presumably independent of Smad activation.(30) Our sequencing results showed that after up-regulating miR-205, the ratio value of Smad4 gene in HeLa cells was 1.3294, which was not statistically significant, while the expression of Smad4 gene in Siha cells was up-regulated. HeLa cells integrate with HPV18 genome. SiHa cells integrated with the HPV16 genome. Therefore, we believe that Smad4 regulates TGF-signaling pathway to exert carcinogenesis and whether it is related to HPV phenotype remains to be explored.

The results of gene microarray in our study reflected the gene expression changes of cervical cancer cells HeLa and SiHa after up-regulating miR-205 or down-regulating *miR-141*.Besides, we analysed the signal pathways in which the DEGs of miR-205 up-regulated SiHa cell participated, but functional experiments were needed to further verify the molecular mechanism of these gene changes. From the comprehensive analysis of microarray, the expression of genes after up-regulating miR-205 changed towards the direction of inhibiting the malignant phenotype of cervical cancer. A group of genes were found, which provided further research direction for illuminating the molecular mechanism of the carcinogenesis of miR-205.

## References

1. Torre LA, Bray F, Siegel RL, Ferlay J, Lortet-Tieulent J, Jemal A. Global cancer statistics, 2012. CA: A Cancer Journal for Clinicians 2015;65(2):87–108.

2. Arbyn M, Castellsague X, de Sanjose S, Bruni L, Saraiya M, Bray F, et al. Worldwide burden of cervical cancer in 2008. Annals of Oncology 2011;22(12):2675–2686.

3. Prasongdee P, Tippayawat P, Limpaiboon T, Leelayuwat C, Wongwattanakul M, Jearanaikoon P. The development of simultaneous measurement of viral load and physical status for human papillomavirus 16 and 18 co-infection using multiplex quantitative polymerase chain reaction. Oncology letters 2018;16(6):6977–6987.

4. Honegger A, Schilling D, Bastian S, Sponagel J, Kuryshev V, Sültmann H, et al. Dependence of Intracellular and Exosomal microRNAs on Viral E6/E7 Oncogene Expression in HPV-positive Tumor Cells. PLOS Pathogens 2015;11(3):e1004712.

5. Liu X, Clements A, Zhao K, Marmorstein R. Structure of the HumanPapillomavirus E7 Oncoprotein and Its Mechanism for Inactivation of the Retinoblastoma Tumor Suppressor. Journal of Biological Chemistry 2005;281(1):578–586.

6. Yu M, Xu Y, Pan L, Feng Y, Luo K, Mu Q, et al. miR-10b Downregulated by DNA Methylation Acts as a Tumor Suppressor in HPV-Positive Cervical Cancer via Targeting Tiam1. Cellular Physiology and Biochemistry 2018;51(4):1763–1777.

7. Peng Y, Croce CM. The role of MicroRNAs in human cancer. Signal Transduction and Targeted Therapy 2016;1(1).

8. Gonzalez-Quintana V, Palma-Berre L, Campos-Parra AD, Lopez-Urrutia E, Peralta-Zaragoza O, Vazquez-Romo R, et al. MicroRNAs are involved in cervical cancer development, progression, clinical outcome and improvement treatment response (Review). Oncol Rep 2016;35(1):3–12.

9. Lu J, Getz G, Miska EA, Alvarez-Saavedra E, Lamb J, Peck D, et al. MicroRNA expression profiles classify human cancers. Nature 2005;435(7043):834–838.

10. Calin GA, Croce CM. MicroRNA signatures in human cancers. Nature Reviews Cancer 2006;6(11):857–866.

11. Xu XW, Li S, Yin F, Qin LL. Expression of miR-205 in renal cell carcinoma and its association with clinicopathological features and prognosis. Eur Rev Med Pharmacol Sci 2018;22(3):662–670.

12. Lu J, Lin Y, Li F, Ye H, Zhou R, Jin Y, et al. MiR-205 suppresses tumor growth, invasion, and epithelial– mesenchymal transition by targeting SEMA4C in hepatocellular carcinoma. The FASEB Journal 2018;32(11):6123–6134.

13. Li L, Li S. miR-205-5p inhibits cell migration and invasion in prostatic carcinoma by targeting ZEB1. Oncology letters 2018;16(2):1715–1721.

14. Chu P, Liang A, Jiang A, Zong L. miR-205 regulates the proliferation and invasion of ovarian cancer cells via suppressing PTEN/SMAD4 expression. Oncol Lett 2018;15(5):7571–7578.

15. Pang H, Yue X. MiR-205 serves as a prognostic factor and suppresses proliferation and invasion by targeting insulin-like growth factor receptor 1 in human cervical cancer. Tumor Biology 2017;39(6):101042831770130.

16. Massague J, Chen YG. Controlling TGF-beta signaling. Genes Dev 2000;14(6):627–44.

17. Yang G, Yang X. Smad4-mediated TGF-beta signaling in tumorigenesis. International journal of biological sciences 2010;6(1):1–8.

18. Shi Y, Massagué J. Mechanisms of TGF-β Signaling from Cell Membrane to the Nucleus. In. United States: Elsevier Inc; 2003. p. 685–700.

19. Baldus SE, Schwarz E, Lohrey C, Zapatka M, Landsberg S, Hahn SA, et al. Smad4 deficiency in cervical carcinoma cells. Oncogene 2005;24(5):810–819.

20. Xu Q, Tong J, Zhang C, Xiao Q, Lin X, Xiao X. miR-27a induced by colon cancer cells in HLECs promotes lymphangiogenesis by targeting SMAD4. PLOS ONE 2017;12(10):e0186718.

21. Chae D, Ban E, Yoo YS, Kim EE, Baik J, Song EJ. MIR-27a regulates the TGF-β signaling pathway by targetingSMAD2 andSMAD4 in lung cancer. Molecular Carcinogenesis 2017;56(8):1992–1998.

22. Du M, Chen W, Zhang W, Tian X, Wang T, Wu J, et al. TGF-β regulates the ERK/MAPK pathway independent of the SMAD pathway by repressing miRNA-124 to increase MALAT1 expression in nasopharyngeal carcinoma. Biomedicine & Pharmacotherapy 2018;99:688–696.

23. Elliott RL, Blobe GC. Role of transforming growth factor Beta in human cancer. J Clin Oncol 2005;23(9):2078–93.

24. Ahmed S, Bradshaw A, Gera S, Dewan M, Xu R. The TGF-β /Smad4 Signaling Pathway in Pancreatic Carcinogenesis and Its Clinical Significance. Journal of Clinical Medicine 2017;6(1):5.

25. Zhu H, Luo H, Shen Z, Hu X, Sun L, Zhu X. Transforming growth factor-β1 in carcinogenesis, progression, and therapy in cervical cancer. Tumor Biology 2016;37(6):7075–7083.

26. Xu Q, Wang S, Xi L, Wu S, Chen G, Zhao Y, et al. Effects of human papillomavirus type 16 E7 protein on the growth of cervical carcinoma cells and immuno-escape through the TGF-β1 signaling pathway. Gynecologic Oncology 2006;101(1):132–139.

27. Lee DK, Kim B, Kim IY, Cho E, Satterwhite DJ, Kim S. The Human Papilloma Virus E7 Oncoprotein Inhibits Transforming Growth Factor-β Signaling by Blocking Binding of the Smad Complex to Its Target Sequence. Journal of Biological Chemistry 2002;277(41):38557–38564.

28. French D, Belleudi F, Mauro MV, Mazzetta F, Raffa S, Fabiano V, et al. Expression of HPV16 E5 down-modulates the TGFbeta signaling pathway. Molecular cancer 2013;12(1):38–38.

29. Peralta-Zaragoza O, Bermudez-Morales V, Gutierrez-Xicotencatl L, Alcocer-Gonzalez J, Recillas-Targa F, Madrid-Marina V. E6 and E7 oncoproteins from human papillomavirus type 16 induce activation of human transforming growth factor beta1 promoter throughout Sp1 recognition sequence. Viral Immunol 2006;19(3):468–80.

30. Maliekal TT, Anto RJ, Karunagaran D. Differential Activation of Smads in HeLa and SiHa Cells That Differ in Their Response to Transforming Growth Factor-β. Journal of Biological Chemistry 2004;279(35):36287–36292.

